# Molecular Basis for ADP-ribose Binding to the Macro-X Domain of SARS-CoV-2 Nsp3

**DOI:** 10.1101/2020.03.31.014639

**Authors:** David N. Frick, Rajdeep S. Virdi, Nemanja Vuksanovic, Narayan Dahal, Nicholas R. Silvaggi

## Abstract

The virus that causes COVID-19, SARS-CoV-2, has a large RNA genome that encodes numerous proteins that might be targets for antiviral drugs. Some of these proteins, such as the RNA-dependent RNA polymers, helicase and main protease, are well conserved between SARS-CoV-2 and the original SARS virus, but several others are not. This study examines one of the proteins encoded by SARS-CoV-2 that is most different, a macrodomain of nonstructural protein 3 (nsp3). Although 26% of the amino acids in this SARS-CoV-2 macrodomain differ from those seen in other coronaviruses, biochemical and structural data reveal that the protein retains the ability to bind ADP-ribose, which is an important characteristic of beta coronaviruses, and potential therapeutic target.

---

The development of antivirals targeting <u>S</u>evere <u>A</u>cute <u>R</u>espiratory <u>S</u>yndrome <u>Co</u>rona<u>v</u>irus <u>2</u> (SARS-CoV-2), the causative agent of the present COVID-19 pandemic,^1^ will most likely focus on viral proteins and enzymes needed for replication.^2^ Like other coronaviruses, SARS-CoV-2 has a large positive sense (+)RNA genome over 30,000 nucleotides long with several open reading frames. Most of the proteins that form the viral replicase are encoded by the “rep 1ab” reading frame, which codes for a 7,096 amino acid-long polyprotein that is ultimately processed into at least 15 functional peptides, five of which are only produced by a translational frameshift event occurring after nsp10 (Fig. 1). Parts of the SARS-CoV-2 rep 1ab polyprotein are very similar to the rep 1ab protein of the coronavirus that caused the SARS epidemic in 2003 (which will be referred to here as SARS-CoV-1), suggesting the that drugs targeting the SARS-CoV-1 nsp5-14 might be effective against SARS-CoV-2. However, some portions of the SARS rep 1ab polyproteins are quite different.

**FIGURE 1.**
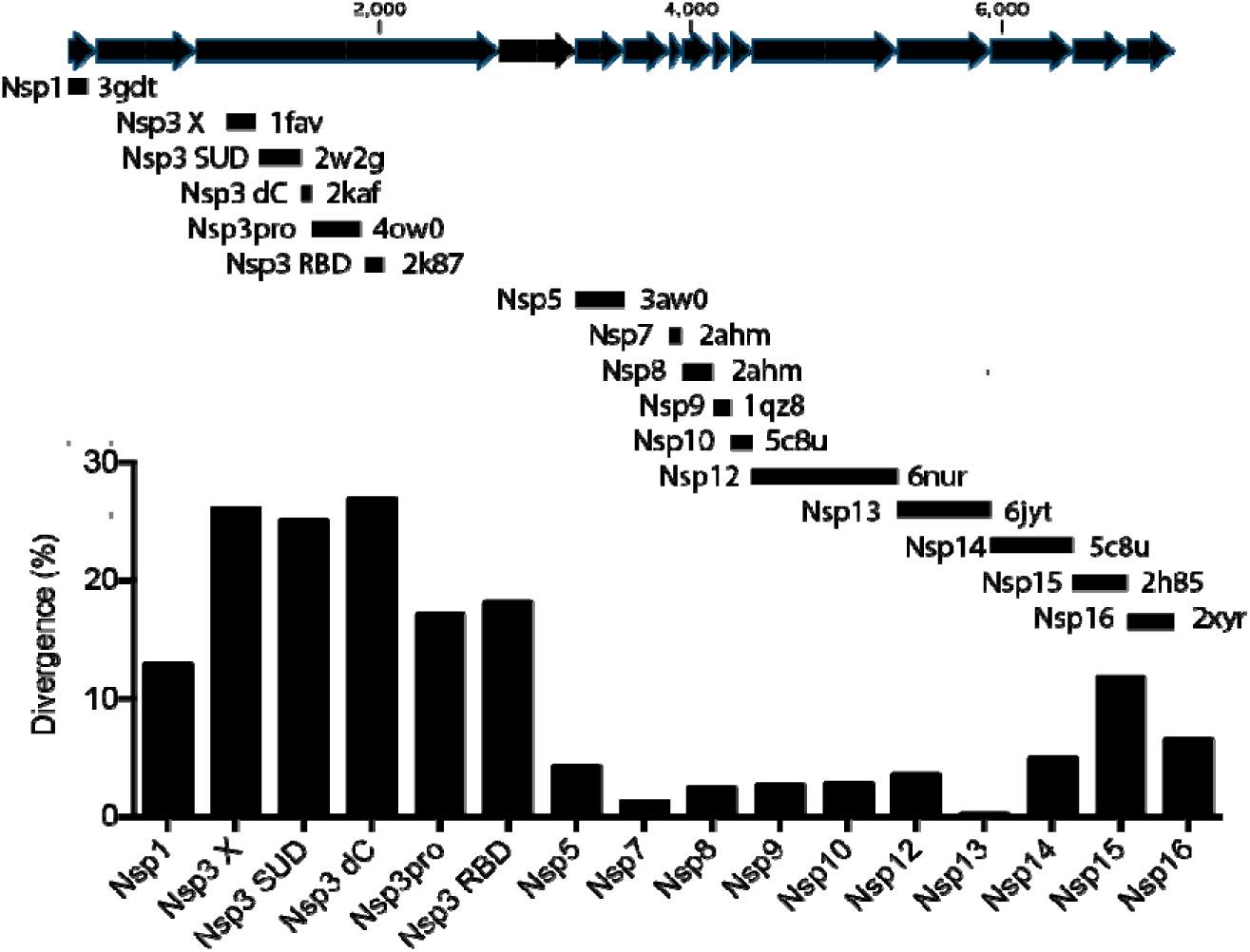
Sequence divergence between potential drug targets in SARS-CoV-1 and SARS-CoV-2. The SARS-CoV-2 rep 1ab peptide sequence was aligned with each of the PDB files listed, which describe an atomic structure of a homologous region of the SARS-CoV-1 rep 1ab polyprotein. Nsp’s are shown in sequence as black arrows (note: there is no “nsp11”, and the translational frameshift occurs after nsp10). The percent of amino that differ in each protein is plotted. Nsp1 is and interferon antagonist.^26^ The Nsp3 X domain is studied here, the Nsp3 SUD consists of tandem macrodomains that bind G-quadruplex structures,^3^ Nsp dC is the C-terminus of the SUD,^27^ Nsp3pro is a papain-like protease,^28^ and Nsp3 RBD is another possible RNA binding domain.^29^ Nsp5 is the main viral protease.^30^ Nsp7 and Nsp8 are polymerase cofactors.^31^ Nsp9 is an RNA binding protein.^32^ Nsp10 is a zinc-binding cofactor for Nsp14 and Nsp16.^33^ Nsp12 is the RNA polymerase.^31^ Nsp13 is a helicase.^34^ Nsp14 is a 3’-5’ exonuclease and a 7-methyltransferase.^33^ Nsp15 is an RNA endonuclease.^35^ Nsp16 is an RNA cap 2’-O-Methyltransferase.^36^

In contrast to the well-conserved SARS nsp5 protease, nsp12 polymerase and nsp13 helicase enzymes, significantly more differences exist between the nsp3 proteins encoded by SARS-CoV-1 and SARS-CoV-2. The most variation occurs in a domain of Nsp3 domain suspected to bind ADP-ribose, that will be referred to here as the “macro X” domain, to differentiate it from the two downstream “SARS unique macrodomains (SUDs),” which do not bind ADP-ribose.^3^ The macro X domain of SARS-CoV-1 is also able to catalyze the hydrolysis of ADP-ribose 1’’ phosphate, albeit at a slow rate.^4^

All the above differences preclude the use of the SARS-CoV-1 macro X domain structures as scaffolds to design compounds that might target this nsp3 region in SARS-CoV-2, especially in light of the observation that the same nsp3 domain from gamma coronaviruses does not bind ADP-ribose *in vitro*.^5^ Because ADP-ribose binding is a property that could be used to identify possible antiviral agents, the ability of the SARS-CoV-2 macro X domain to bind ADP-ribose was examined using a recombinant purified protein and isothermal titration calorimetry (ITC). We also determined the structure of the SARS-CoV-2 macro X domain in order to examine the biochemical context of ADP-ribose binding and to provide data for rational inhibitor design or *in silico* screening.

## MATERIAL & METHODS

### Gene Synthesis

To facilitate comparison between SARS-CoV-1 and SARS-CoV-2, a protein expression vector was generated similar to the one used by Eglott *et al.*^6^ To this end, a codon optimized open reading frame was synthesized by GenSript (Piscataway, NJ) that encodes the Macro X domain with an N-terminal TEV-cleavage site flanked by *Nhe*I and *Bam*H1 restriction sites. This open reading frame was cloned into pET21b to give plasmid pET21-COVID-MacroX. The pET11-COVID-MacroX plasmid was used to transform BL21(DE3) cells.

### Protein Purification

Colonies of BL21(DE3) cells harboring the pET21-COVID-MacroX plasmid were used to inoculate 3 ml of LB medium containing 100 mg/ml ampicillin. The starter culture was incubated at 37 °C with shaking at 225 rpm. After the cells grew to an OD_600_ of 1.0, they were transferred to 1 liter of fresh medium containing ampicillin. After the cells reached an OD_600_ of 1.0 again, protein production was induced with 1 mM isopropyl-β-D-thiogalactoside. After growing 16 h at 23 °C, the cells were harvested by centrifugation at 4,000 rpm, 4°C. The resulting cell pellet was suspended in 25 mL of IMAC buffer (20 mM Tris pH 8, 0.5 M NaCl), sonicated on ice for five 1 min bursts, with 2 min rests between, and clarified by centrifugation at 10,000g for 30 min. The supernatant was loaded onto a 5 ml Ni-NTA column and the fractions were eluted with a step gradient from 5 to 500 mM imidazole. Fractions containing the macro X domain protein (5 ml total) were loaded on a 250 ml Sephacryl S300 gel filtration column and eluted with 10 mM MOPS, 150 mM NaCl. Concentration of the purified protein was determined by measuring absorbance at 260 nm using a molar extinction coefficient of 10,555 M^-1^ cm^-1^, which was calculated with the ProtParam tool (https://web.expasy.org/protparam/).

### Isothermal Titration Calorimetry (ITC)

Binding of ADP-ribose to the SARS-CoV-2 macro X domain was measured using a Nano ITC (TA Instruments). Before starting the measurement, samples of both ligand and protein were diluted in 10 mM MOPS, 150 mM NaCl (pH 7) and were degassed at 400 mmHg for 30 minutes. Measurements were taken at 20 °C by injecting 2.0 µl aliquots of 500 µM ADP-Ribose (Sigma) to 50 µM protein (175 µl initial volume) with 250 rpm stirring rate. Using NanoAnalyze Software (v. 3.11.0), data were fitted by non-linear regression to an independent binding model. Briefly, after baseline correction, background heats from ligand-to-buffer titrations were subtracted, and the corrected heats from the binding reaction were used to find best fit parameters for the stoichiometry of the binding (n), free energy of binding (ΔG), apparent enthalpy of binding (ΔH), and entropy change (ΔS). Dissociation constants (K_d_) were calculated from the ΔG.

### Crystallization and Structure Determination

In preparation for crystallization experiments, the purified SARS-CoV-2 macro X domain protein was cleaved with tobacco etch virus (TEV) protease to remove the Nterminal His_6_-tag and passed back through the Ni-NTA column. The flow-through fractions were desalted into 10 mM HEPES, pH 7.2 using a 2×5ml HiTrap desalting column (GE Life Sciences) and concentrated to 10 mg/mL in a centrifugal concentrator. This preparation of the protein was mixed 1 μL:1 μL with the Morpheus HT screen reagents (Molecular Dimensions) in a 96-well SwissSci MRC UV-transmissible sitting drop plate. Large, diffraction-quality crystals grew directly from a number of the screen conditions. The crystal ultimately used for structure determination grew from condition D9: 0.12 M alcohols [0.02 M each 1,6-hexanediol, 1-butanol, 1,2-propanediol, 2-propanol, 1,4-butanediol, and 1,3-propanediol], 0.1 M buffer system 3, pH 8.5 [0.05 M each TRIS and bicine], and 30% precipitant mix 1 [20% poly(ethylene glycol) (PEG) 500 monomethylether, 10% PEG 20,000]. Large, thick plates grew within 1 week at 22 °C. Owing to the high concentration of PEG 500 MME, the crystal did not require additional cryo-protection and was flash-cooled by looping it directly from the sitting drop and plunging it into liquid nitrogen.

Diffraction data were collected on Life Sciences Collaborative Access Team (LS-CAT) beamline 21-ID-F at the Advanced Photon Source of Argonne National Laboratory. The wavelength at this station is fixed at 0.9787 Å; the detector is a MarMosaic M300 charge-coupled device. The data were collected with an oscillation width of 0.5° per image for a total oscillation of 180°. The data were indexed and integrated with DIALS ^7, 8^ as implemented in version 7.2 of the CCP4 software suite.^9, 10^ Data scaling and reduction was performed using AIMLESS.^11–13^ Data collection statistics are provided in Table 1.

**Table 1.** Crystallographic data collection and model refinement statistics for the SARS-CoV-2 Macro X domain.

| Data collection |  |
| --- | --- |
| Resolution range (Å) (last shell)<sup>a</sup> | 43.32 – 0.95 (0.97 – 0.95) |
| Space Group | P 2<sub>1</sub> 2<sub>1</sub> 2<sub>1</sub> |
| *a*, *b*, *c* (Å) | 43.3, 54.4, 67.6 |
| $\alpha$, $\beta$, $\gamma$ (°) | 90.0, 90.0, 90.0 |
| R<sub>merge</sub><sup>a</sup> | 0.063 (0.348) |
| R<sub>meas</sub><sup>a</sup> | 0.075 (0.475) |
| R<sub>pim</sub><sup>a</sup> | 0.039 (0.320) |
| CC<sub>1/2</sub><sup>a</sup> | 99.5 (89.1) |
| No. of unique reflections<sup>a</sup> | 99442 (4199) |
| Completeness (%)<sup>a</sup> | 98.4 (85.0) |
| Multiplicity<sup>a</sup> | 6.4 (2.6) |
| $\langle I/\sigma(I) \rangle$<sup>a</sup> | 13.1 (1.5) |
| Model Refinement |  |
| Reflections used in refinement | 99335 (2606)<sup>b</sup> |
| Reflections used for R<sub>free</sub> | 5046 (125) |
| R<sub>cryst</sub> (R<sub>free</sub>) | 0.119 (0.137) |
| Wilson B-factor (Å<sup>2</sup>) | 7.4 |
| Average B factor (Å<sup>2</sup>) | 13.9 |
| Protein atoms | 10.2 |
| Solvent | 28.2 |
| Root-mean-square (RMS) deviations |  |
| Bond lengths (Å) | 0.010 |
| Bond angles (°) | 1.16 |
| Coordinate error (Å) | 0.06 |
| Ramachandran statistics |  |
| Favored/allowed/outliers (%) | 98.2/1.8/0.0 |
| Rotamer outliers (%) | 1.2 |
| Clashscore | 1.70 |

<sup>a</sup>Values in parentheses apply to the high-resolution shell indicated in the resolution row<sup>b</sup>The limits of the high-resolution bin for refinement were 0.96 – 0.95Å

The structure was determined by molecular replacement in PHASER^14^ using the model of the SARS-CoV-1 macro X domain as the search model (PDB ID 2FAV^6^). The model underwent iterative rounds of (re)building in COOT^15^ and refinement in PHENIX.refine.^16, 17^ The very high resolution of the data justified a full anisotropic treatment of the protein and solvent temperature factors. Model refinement and validation statistics are provided in Table 1. The coordinates were deposited in the Protein Data Bank with accession code 6WEY.

## RESULTS AND DISCUSSION

### Hypervariability in the nsp3 Macro X domain

The structures of most of the soluble portions of the SARS-CoV-1 nsp proteins have been examined at atomic resolution to help understand coronavirus replication and facilitate antiviral drug discovery. The amino acid sequences of each of these proteins were compared with the homologous regions of the rep 1ab protein encoded by SARS-CoV-2 (GenPept Accession YP_009724389). The most similar proteins were the RNA helicases (nsp13) which are identical in all but one of their 603 amino acids: a conservative Val to Ile substitution near their C-termini). The RNA-dependent RNA polymerases (nsp12) are also well-conserved, sharing all but 34 of 955 amino acids. The primary protease that cleaves the polyprotein (nsp5) is also similar in SARS-CoV-1 and SARS-CoV-2, with only 13 amino acids that differ among 306 (4.2% different) (Fig. 1).

At the other end of the spectrum are the nsp3 proteins, which are notably more different in the two SARS viruses. Nsp3 is a large multidomain membrane-bound protein,^18^ and its clearest role in viral replication is cleaving the rep polyprotein. Over 17% of the amino acids in the nsp3 protease domain differ between SARS-CoV-1 and SARS-CoV-2. Other parts of nsp3 are even more variable, like the macrodomains that lie N-terminal to the nsp3 protease domain. Macrodomains consist of four helices that surround a mixed beta sheet. A ligand-binding pocket that typically binds ADP-ribose or related compounds lies between the helices and the sheet.^19^ SARS-CoV-1 has three macrodomains in tandem, but only the first binds ADP ribose. The amino acid sequences of this macro X domain differ by 26% between SARS-CoV-1 and SARS-CoV-2 (Fig. 1).

Many of the 47 variant residues in the 180 amino acid long SARS macro X domain are clustered near its N-terminus in a region that is particularly variable in the three *Coronaviridae* genera (Fig. 2). Macro X domains from coronaviruses that cause the common cold (alpha coronaviruses)^5^ and the beta coronaviruses like SARS-CoV-1 and Middle East respiratory syndrome coronavirus (MERS-CoV)^20^ all bind ADP ribose. However, the same domain in gamma coronaviruses does not bind ADP-ribose.^5^

**FIGURE 2.**
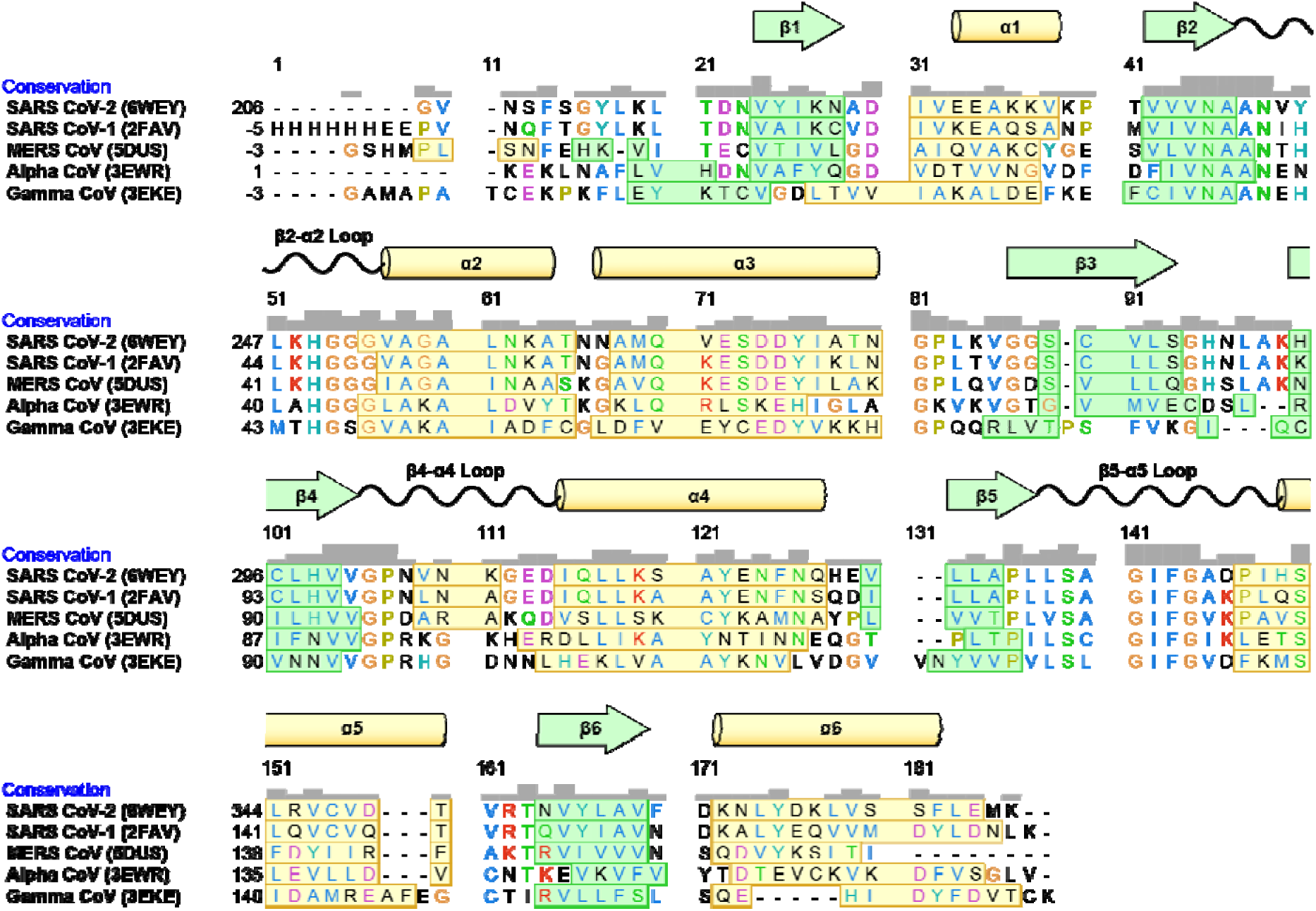
Variation in the macro X domains of coronaviruses. (A) Macro X domain structures were aligned using the “MatchMaker” function of UCSF Chimera (v. 1.14).^37^ Amino acids are colored by class with beta sheets in green boxes and alpha helices in yellow boxes.

### Expression and Purification of the SARS-CoV-2 Macro X domain

An *E. coli* expression vector for the macro X domain was built to express and N-terminally Histagged protein similar to a SARS-CoV-1 protein studied by Egloff *et al.*^6^ Upon induction, a one liter culture of BL21(DE3) cells harboring the vector expresses 50-100 mg of the macro X domain protein that can be purified in one step to apparent homogeneity using immobilized metal affinity chromatography (Fig. 3A). The protein was polished further with gel filtration chromatography and concentrated before analysis and crystallization.

**FIGURE 3.**
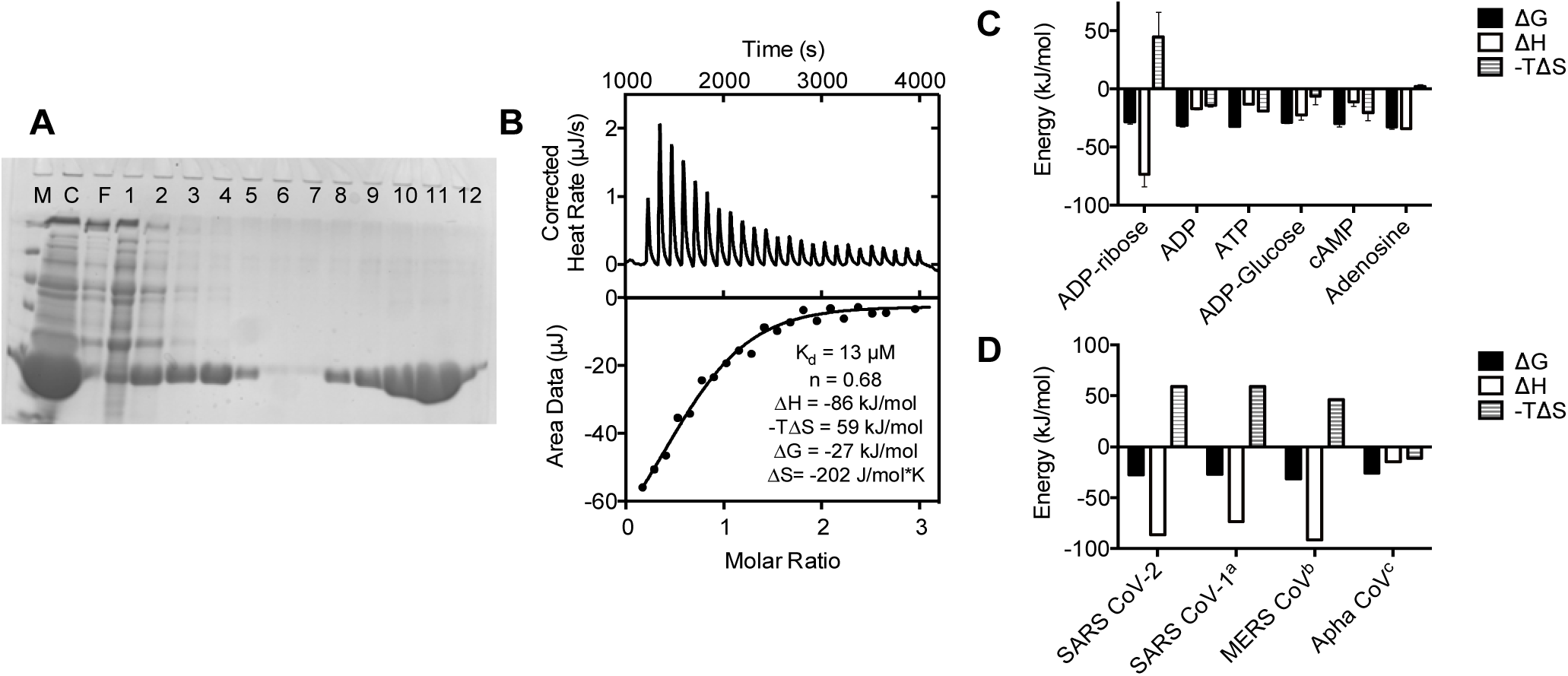
The SARS-CoV-2 macro X domain binds ADP ribose. (A) 15% SDS PAGE showing 10 µL samples of: a soluble crude lysate of induced BL21(DE3) cells harboring the plasmid p21-COVID-MacroX (lane C), proteins the do not bind a Ni-NTA column (F), and fractions eluted from a Ni-nitrilotriacetic acid column during an imidazole step gradient from 0 mM (lanes 1-3), 5 mM (lanes 4-6), 40 mM (lanes 7-9), and 500 mM (lanes 10-12). Protein markers (lane M) are 116, 66.2, 45, 35, and 25 kDa. (B) Example ITC experiment in which purified SARS-CoV-2 nsp3 macro domain was titrated with ADP ribose. (C) ITC experiments like those shown in panel B were repeated three times with each of the compounds listed. Means are plotted and error bars are standard deviations. Average (±SD) dissociation constants were 10 ± 4 µM for ADP-ribose, 8 ± 9 µM for ADP, 3 ± 3 µM for ATP, 6± 4 µM for ADP-glucose, 2 ± 1 µM for cAMP, and 2 ± 1 µM for adenosine. (D) Comparison of the thermodynamics of ADP-ribose binding by macro X domains from SARS-CoV-2 (data from panel C), SARS-CoV-1, MERS-CoV, and an alpha coronavirus. ^a^Data from Egloff et al.^32^ ^b^Data from Cho et al.^20^ ^c^Data from Piotrowski et al.^5^

### Nucleotide Binding by the SARS-CoV-2 Macro X domain

Repeated ITC experiments revealed that the purified recombinant protein bound ADP-ribose (Fig. 3C) with a dissociation constant of 10 ± 4 µM (uncertainty is the standard deviation of K_d_’s from independent titrations). To examine binding specificity, similar titrations were repeated with related nucleotides. The SARS-CoV-2 protein bound ADP, cAMP, ATP and ADP-glucose (Fig. 3D). All nucleotides lacking the ribose moiety bound with similar high affinities, but none bound with an enthalpy change like that seen with ADP-ribose, suggesting specific contacts form between ADP ribose the SARS-CoV-2 protein. Based on what is seen in the SARS-CoV-2 structures below, these contacts likely occur with the conserved D226 and N244 (positions 30 and 43 in the numbering above the alignment in Fig. 2). None of the other nucleotides bind with an entropic penalty as seen with ADP-ribose, either, suggesting that the ribose moiety becomes structured when bound to the macrodomain.

The energetics of ADP-ribose binding to the SARS-CoV protein are similar to those seen with the same protein from SARS-CoV-1,^6^ MERS-CoV.^20^ Enthalpy and entropy of binding were also very similar for all three protein (Fig. 3D). Unlike what is seen with the macro X protein from an alpha coronavirus,^5^ enthalpy appears to drive ADP-ribose binding to the macro X domains of the three beta coronaviruses.

### Structure of the SARS-CoV-2 Macro X Domain

The SARS-CoV-2 macro X domain (NSP3 residues 207 to 277) crystallized in space group P2_1_2_1_2_1_ with 1 molecule per asymmetric unit. These crystals had a solvent content of 43% and diffracted extremely well. The final resolution limit of the data was set at 0.95 Å (Table 1). The quality of the electron density maps is correspondingly excellent (Fig. 4A). The section of the structure depicted in this image is located on the surface of the protein and the B-factors of these residues are close to the average B-factor of the protein (7.2 vs 10.2 Å^2^), indicating that this sample accurately represents the overall quality of the maps. The final model contains the entire sequence from V207 to S377 of nsp3, an N-terminal glycine residue that was left from the TEV-protease cleavage, and 374 solvent molecules. The R_cryst_ and R_free_ values of the final model were 0.119 and 0.137, respectively (Table 1).

**FIGURE 4.**
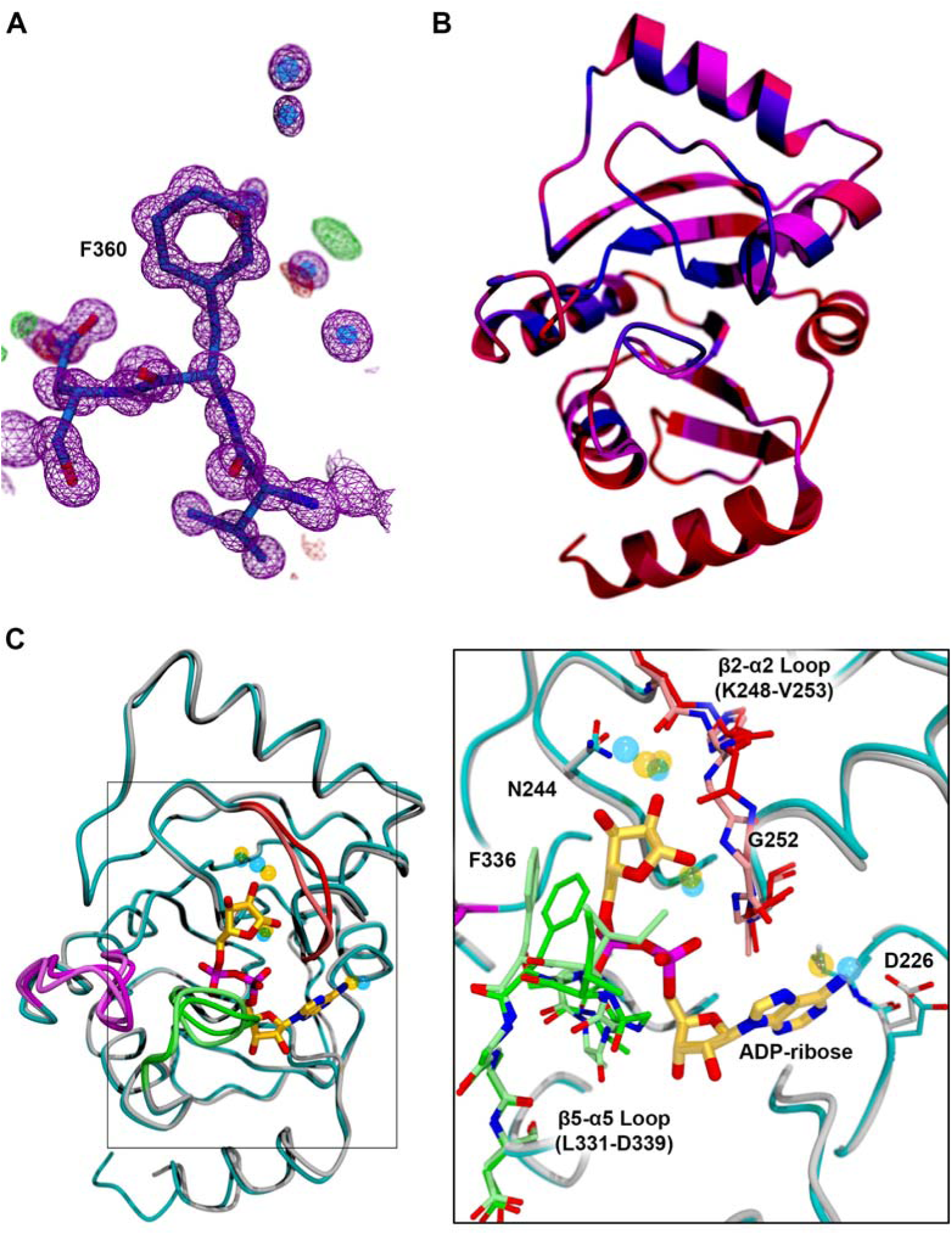
The SARS-CoV-2 macro X domain structure. (A) The electron density is shown for a representative portion of the structure (residues 359-361) on the surface of the protein. The 2mFo-DFc map is contoured at 1.5σ and is shown as a magenta mesh. The mFo-DFc (difference) maps are shown at + and – 3.0σ as green and red mesh, respectively. (B) Ribbon diagram of the SARS-CoV-2 macro X domain structure colored according to the sequence conservation plot in Fig. 2 as a gradient from red (low conservation; <10%) to blue (highly conserved; 100%) through magenta. As can be seen in the sequence alignment, the N- and C-termini are particularly poorly conserved. (C) Overlay of the structure of the SARS-CoV-2 macro X domain bound to ADPribose determined by Michalska et al. of the CGSID (PDB ID 6W02) with the ultra-high-resolution structure of the unliganded protein determined here. ADP-ribose is shown in ball-and-stick with the carbon atoms colored gold. The backbone trace of the unliganded structure is colored cyan, and that of the ADP-ribose-bound model is colored grey. There are three loops with significantly different conformations in the two structures. In the unliganded structure, the β2-α2 loop is colored bright red, the β4-α4 loop is colored purple, and the β5-α5 loop is colored bright green. The same regions of the ADP-ribose-bound structure are colored pale red, pale purple, and pale green, respectively. The transparent blue and yellow spheres represent water molecules bound to the unliganded (transparent blue) and ADP-ribose-bound (transparent yellow) forms of the protein. Interestingly, several of the water molecules interacting with ADP-ribose in 6W02 can also be found in the unliganded structure of the protein. The inset shows a close-up of the boxed region colored according to the same scheme. The β2-α2 and β5-α5 loops, which contact ADPribose, are shown in ball-and-stick. Note that the β2-α2 loop rotates ~180° to allow it to make a hydrogen-bonding interaction with the 1’-hydroxyl of the ribose moiety. Also, the phenylalanine residue in the β5-α5 loop (F336) would clash with the β-phosphate and ribose of ADP-ribose if the β5-α5 loop did not adopt a different conformation.

The tertiary structure, as expected, ranges from nearly identical to very similar to those of other coronavirus macro domains, including: SARS CoV-1 (2FAV^6^; 74.7% sequence identity) with a root mean square deviation (RMSD) value for 162 of 172 Cα atoms of 0.6 Å; MERS CoV (5DUS^20^; 42.2% identical) with a 1.2 Å RMSD for 161 of 172 Cα atoms; human alpha coronavirus 229E (3EWR^21^; 32.5% identical) with a 1.5 Å RMSD for 154 of 172 Cα atoms; feline coronavirus (FCoV; 3JZT^22^ 26.8% identical) with a 1.5 Å RMSD for 153 of 172 Cα atoms; and the gamma CoV infectious bronchitis virus (IBV; 3EWP^21^ 26.7% identical) with a 2.1 Å RMSD for 150 of 172 Cα atoms. The regions of high sequence conservation are not clustered in any particular region(s) of the molecule (Fig. 4B). This makes sense in light of the fact that the protein atoms involved in hydrogen bonding interactions with the ligands in these structures are more often part of the main chain; there are relatively few interactions of side chains with the ligands.

At the time of writing, we discovered that Michalska *et al.* of the Center for Structural Genomics of Infectious Diseases (CSGID) deposited coordinates for a very similar construct of the SARS-CoV-2 macro X domain including from E206 to E275 of the nsp3 protein, plus an additional 4 residues at the N-terminus (6VSX; unpublished). Their crystals also allowed binding of ADPribose (6W02) and AMP (6W6Y), while ours seemed to be packed too tightly to permit ligands to access the binding site (data not shown). We compared our ultra-high-resolution model of the unliganded protein to the ADP-ribose-bound form. The RMSD values for the fitting, done by secondary structure matching (SSM)^23^ as implemented in COOT, is 0.59 Å for 165 of 172 Cα atoms. This is very similar to the RMSD values of the free protein (6VXS; 0.66 Å) and the AMP-bound form (6W6Y; 0.50 Å), indicating that there are no large conformational changes that occur upon ligand binding. In fact, the only notable conformational changes occur in three surface-exposed loops in or near the ligand-binding pocket (Fig. 4C). These loops connect strand β2 with helix α2 (the β2-α2 loop), strand β4 with helix α4 (β4-α4 loop), and strand β5 with helix α5 (β5-α5 loop). The subtle change in conformation of the β4-α4 loop (purple in Fig. 4C) appears to be the result of crystal contacts and not the direct influence of ADP-ribose binding. The other two loops are more intimately involved in ligand binding. The main chain of the α2-β2 loop rotates 180° in order to allow the amide N atom of G252 to participate in a hydrogen bonding interaction with the 1’-hydroxyl of the ribose moiety of ADPribose. This loop also carries N244, which directly interacts with the ribose. The phenyl ring of F336 in the β5-α5 loop occupies the portion of the binding pocket in the unliganded structure that is occupied by the β-phosphate of ADP-ribose. Thus, without a rearrangement of the β5-α5 loop, ADP-ribose would not be able to bind.

### Conclusion

The significance of the study stems mainly from the demonstration that the SARS-CoV-2 macro X domain binds ADP ribose. This is the first step needed to justify screens for potential antivirals that bind in place of ADP ribose. More work needs to be done, however, to understand the antiviral potential of such compounds because the biological role for ADP-ribose binding is still not fully understood. Some work with alpha coronaviruses suggest that ADP-ribose binding by the macro X domain is not needed for viral replication.^24^ However, studies with other (+)RNA viruses suggest that macrodomains are essential for virulence.^25^ This work is also noteworthy because the synthetic codon-optimized plasmid reported here produces up to 100 mg of soluble macro X domain protein per liter of *E. coli* culture, and this protein retains a high affinity for ADP ribose. The protein could be used for structural studies and screening campaigns. Screening assays with the SARS-CoV-2 protein might actually be more efficient, since the SARS-CoV-2 protein binds ADP-ribose somewhat more tightly (K_d_ = 10 µM) than the SARS-CoV-1 protein (K_d_ = 24 µM). The recombinant protein reported here, together with the detailed structural information, might also be useful to others developing SARS-CoV-2 diagnostics and/or therapeutics.

## Accession Codes

SARS-CoV-2 Rep 1ab YP_009724389 (NCBI) SARS-CoV-2 Macro X 6WEY (PDB)

## ACKNOWLEDGMENTS

The authors are grateful to Matt McCarty, Garrett Breit, Hayden Aristizabal, Trevor R. Melkonian, Dante A. Serrano and Prof. Ionel Popa for valuable technical assistance and helpful discussions.

This research used resources of the Advanced Photon Source, a U.S. Department of Energy (DOE) Office of Science User Facility operated for the DOE Office of Science by Argonne National Laboratory under Contract No. DE-AC02-06CH11357. Use of the LS-CAT Sector 21 was supported by the Michigan Economic Development Corporation and the Michigan Technology Tri-Corridor (Grant 085P1000817).

## Insert Table of Contents artwork

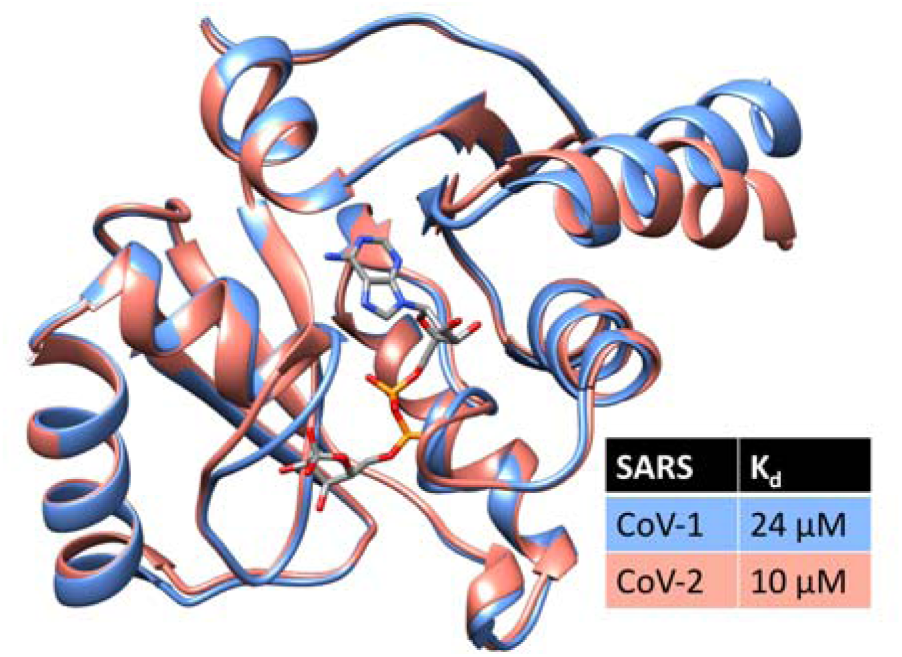

